# Motor Cortex Encodes A Temporal Difference Reinforcement Learning Process

**DOI:** 10.1101/257337

**Authors:** Venkata S Aditya Tarigoppula, John S Choi, John P Hessburg, David B McNiel, Brandi T Marsh, Joseph T Francis

## Abstract

Temporal difference reinforcement learning (TDRL) accurately models associative learning observed in animals, where they learn to associate outcome predicting environmental states, termed conditioned stimuli (CS), with the value of outcomes, such as rewards, termed unconditioned stimuli (US). A component of TDRL is the value function, which captures the expected cumulative future reward from a given state. The value function can be modified by changes in the animal’s knowledge, such as by the predictability of its environment. Here we show that primary motor cortical (M1) neurodynamics reflect a TD learning process, encoding a state value function and reward prediction error in line with TDRL. M1 responds to the delivery of reward, and shifts its value related response earlier in a trial, becoming predictive of an expected reward, when reward is predictable due to a CS. This is observed in tasks performed manually or observed passively, as well as in tasks without an explicit CS predicting reward, but simply with a predictable temporal structure, that is a predictable environment. M1 also encodes the expected reward value associated with a set of CS in a multiple reward level CS-US task. Here we extend the Microstimulus TDRL model, reported to accurately capture RL related dopaminergic activity, to account for M1 reward related neural activity in a multitude of tasks.

**Significance statement:** There is a great deal of agreement between aspects of temporal difference reinforcement learning (TDRL) models and neural activity in dopaminergic brain centers. Dopamine is know to be necessary for sensorimotor learning induced synaptic plasticity in the motor cortex (M1), and thus one might expect to see the hallmarks of TDRL in M1, which we show here in the form of a state value function and reward prediction error during. We see these hallmarks even when a conditioned stimulus is not available, but the environment is predictable, during manual tasks with agency, as well as observational tasks without agency. This information has implications towards autonomously updating brain machine interfaces as others and we have proposed and published on.

## Introduction

When learning to ride a bike, we learn through trial-and-error, with delays between actions and consequences, such as rewards like staying upright on the bike, or punishments like falling. Trial-and-error learning and forming passive stimulus-outcome associations are well modeled by reinforcement learning (RL) (1-5). When RL is used by a learning agent toward optimal control it relies on “reward”, which can be positive or negative, as feedback, where the goal of the agent is to accumulate the maximum amount of temporally discounted reward (6). In most scenarios, reward outcome may be subject to delays. These delays make it difficult to determine how to assign credit to actions, or states, that have come before a reward. This is called the credit assignment problem. Temporal Difference (TD) learning methods can address this problem. Under TDRL, the agent learns the expected temporally discounted reward (value function) for each of the states visited leading to reward. The agent can then utilize these learned estimates of the state value function at a given state to select an appropriate action to maximize reward (6,7). This same TDRL learning machinery can be used even when not selecting actions, such as during classical conditioning paradigms.

Phasic neural activity in dopaminergic brain centers is similar to the TD error signal, a specific form of reward prediction error, which we will simply refer to as RPE in this work (RPE, 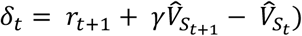, which is the difference between the current state’s estimated value 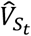, the temporally discounted estimated value of the next state 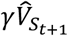 and the immediate reward *r*_*t*+1_ obtained (8,9,10). In this equation *S*_*t*_ is the state, such as the conditioned and unconditioned stimuli in the task at time *t*, and *γ* factor. Note that some of the visual aspects of the task may take part in the brain’s actual state representation, but we are only including explicit stimuli and the time within a trial as part of our state representation to be modeled below, as we show this state representation to be sufficient for this work.

Dopamine has been shown necessary for long-term potentiation in the motor cortex associated with sensorimotor learning (11,12), possibly bridging TDRL theory with sensorimotor learning. Tonic dopaminergic activity has been shown to act like a value function, in this regard; dopamine can “charge” the nervous system, acting as a motivational signal (13,14). Thus, dopamine could have at least two influences on the motor cortex, one gating synaptic plasticity toward sensorimotor learning, and the other “charging” neural activity, possibly priming the system for action.

Previously we, and others, have shown that reward modulates single units and local field potentials in the primary sensorimotor cortices (M1, S1) of non-human primates (2,15-17) and frontal regions influencing M1 (18). We have proposed that neural correlates of reward could be used towards an autonomously updating brain machine interface (2,17). Here we expand on this previous work, showing that M1 activity displays all the hallmarks of a temporal difference reinforcement learning (TDRL) process. We show that M1 activity responds to both expected and unexpected reward delivery. M1 acquires a response to a conditioned stimulus (CS) following conditioning when the CS is used to cue reward. Reward prediction error, an important aspect of the TDRL process, is also encoded in M1 (15) and after an unexpected reward omission. If reward is temporally predictable, then M1 tracks the underlying temporal structure as well. These activity patterns are seen during both manual and observational tasks bilaterally in M1. In addition, if multiple levels of reward are used, with associated conditioned stimuli, M1 activity encodes the expected value following CS presentation in a linear fashion, similar to that reported in the striatum (19).

## Results

We utilized two tasks for our work as seen in figure 1, to probe and understand the M1 neural dynamics in response to explicit/implicit reward expectations and prediction errors. The first was a cued center out reaching task (CCT) where the non-human primates (NHPs) made reaching movements to a single color cued target (Fig.1.a). The color indicated the value of the target, in CCT the value was either rewarding (R, red) or non-rewarding (NR, blue). NHPs A and Z, proficient in performing the CCT, were implanted in the contralateral and ipsilateral M1 respectively. NHPs S and PG, implanted in their contralateral M1, performed a grip force (GF) task and its variations (Fig.1.b). These NHPs were trained to apply an instructed amount of grip force while a virtual cylinder was automatically transported to a target location. In the cued GF task the NHPs were cued (number of green squares) at the beginning of a trial as to the reward they could expect on a successfully completed trial. The uncued GF task had no explicit CS. A reward omission GF task was also performed to test for reward prediction error in M1. We conducted both operant conditioning versions of the CCT and GF tasks, which required manual reaching movements, as well as classical conditioning versions were trials were observational, and the NHP simply watched the task play out before them, while still receiving the outcome value from the task. Sessions in CCT or GF tasks with random sequences of R and NR trials were termed chance trial value predictability (TVP), whereas sessions where R and NR alternated were termed complete TVP sessions. Further details on the behavioral tasks are available in the Methods section.

**Figure 1:**
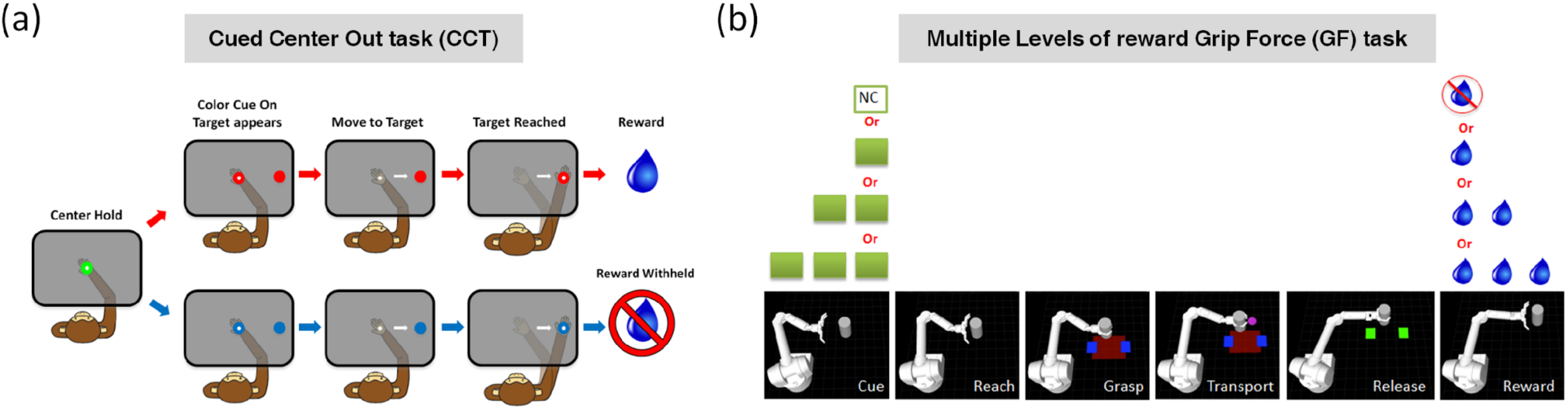
Behavioral tasks. (a) Manual and observational (OT) versions of a Cued Center Out Reaching task (CCT) (*2*). The manual task required a reaching movement from center start position to a single peripheral target. Trial values were cued via the color of the reaching target as either rewarding (red), or non-rewarding (blue) if successful. If a non-rewarding trial was unsuccessful the subject had to repeat it until successful in order to move on to a possibly rewarding trial. Observational CCT required the NHP to simply watch as the task played out with constant speed cursor movements to the target cued as rewarding or non-rewarding. Trial sequence values could be either random or predictable. (b) Cued manual and observational Grip Force (GF) task (see methods). Briefly, the subject watched a projected simulation of an anthropomorphic robotic arm and hand making reaching movements to a target cylinder, at which time the subject had to control the grip force, cued visually by the width of the red rectangle, via a grasped force transducer in order for the robot to grasp the object and automatically transport it to a target location, as long as the subject continued to produce the target grip-force, cued via the blue rectangles, and release the force when the robot reached the target zone. The conditioned stimulus (CS, green squares shown to the subject) cued the NHP of the value; each green square represents one reward volume of juice.

### M1 units have heterogeneous representations of conditioned and unconditioned stimuli

Multiple units in M1 have been shown to respond to the conditioned (CS) and unconditioned stimuli (US) (2,15-17). We have also reported previously (2) and in Fig.3 that the average activity of M1 is significantly different throughout a rewarding trial with respect to a non-rewarding trial. Interestingly, we observed that individual M1 units did not necessarily mimic a behavior similar to the average M1 activity. Some of the M1 units (unit 6, Fig.2) were active only around the reward predicting CS, whereas others (unit 1, 3, 4, Fig.2) were active only around the reward delivery period (US). Some were active around both the CS and US (unit 2, Fig.2). We also found units that showed a negative modulation following the US (unit 3, Fig.2), which increased its difference in activity between R and NR trials as the NHP progressed to later sessions on the same day, possibly due to continued learning of the CS-US.

**Figure 2:**
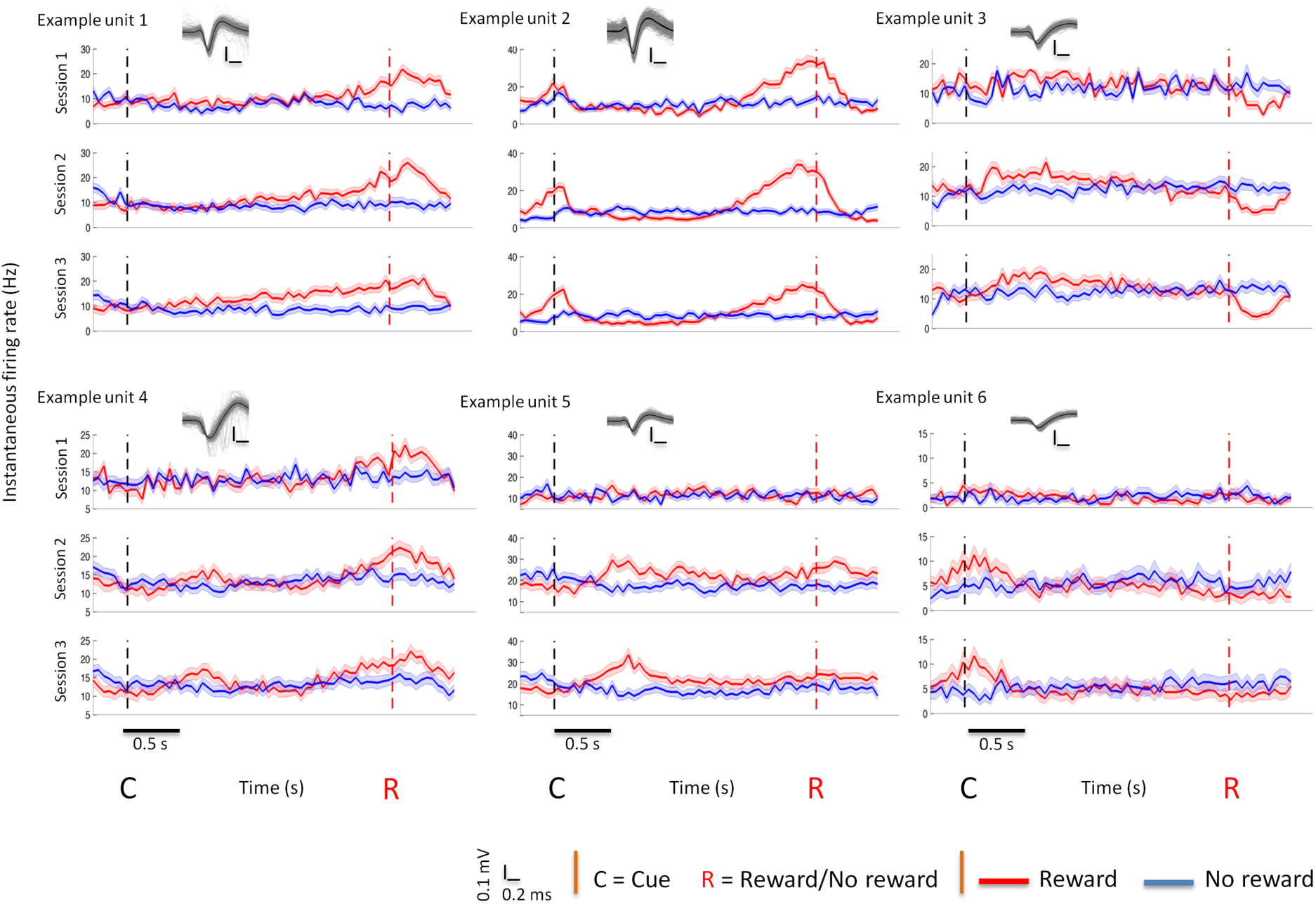
Average activity of example units from NHP A across R and NR trials for each of three CCT observation sessions in a single day. 100 example unit waveforms (grey lines) along with the mean unit waveform (dark black line) for each example unit are also presented. Neural data was binned at 50ms. The vertical dashed black line represents the time of the cue, and the vertical dashed red line represents the average time to reward delivery. Units 1 and 4 show an increased average activity in rewarding trials around the reward delivery period. Unit 2 shows an increased average activity in rewarding trials around both the reward-predicting cue and reward delivery periods. Unit 3 decreased its activity in rewarding trials immediately after reward delivery was initiated. Units 5 and 6 increased their activity either at the reward prediction cue presentation or after the cue presentation but prior to the reward delivery period.

### M1 encodes a state value function

In Fig.3, we present results from the CCT task. In Fig.3.a we have plotted results from the observational (OT) CCT task (Fig.1.a), which is a classical conditioning paradigm, for an example single unit from contralateral M1 that displayed an activity pattern consistent with the evolution of a state value function while learning a cue (CS) - reward (US) association. We have plotted the unit’s peri-cue-time-histogram (PCTH) for R and NR trials, for three sessions, broken up into three equal parts each (Fig.3.a.1 - a.9). As this data comes from the observational (OT) version of the task, confounds of movement related activity are reduced. In session 1 the only time bins with significant differences between R and NR trials are post reward (Fig.3.a1 - a.3), and as experience is gained, from sessions 2 - 3, and putative learning of the association builds, there is movement of significant differences between R and NR trials propagating backward in time towards the presentation of the CS. There was an increase in the number of bins that showed significant differences in the median activity between R and NR trials, including the large increase that occurred after a short break between session 1 and 2 (Fig.3.a3 vs. a4). At the end of these three sessions one can see a peak of activity post CS and before/at reward delivery (Fig.3.a.7-9). We have plotted the average neural activity for a subpopulation of single/multi units that correlated with reward value, separately for R and NR trials, in red and blue respectively (Fig.3.b.2-6). Specifically, this subpopulation is the average of the top 10% of the full population after rank ordering with respect to individual unit’s correlation with reward. In both the OT Fig.3.b, and manual, Fig.3.c, versions of the CCT task, we see that the difference between R and NR trials, grows earlier in the trials as the subject continues the task over time, as in Fig.3.a for the single unit.

We have plotted the PCTH for R and NR trials for all units as false color plots in Fig.3.b, c.2-6. In each false color subplot, the black line is the mean of the full population. Learning also took place for the population mean in the OT-CCT where the trial sequence was completely predictable with R-trials always followed by NR-trials, and repeating. Thus, this was a very stable and predictable environment. Notice that in the OT-CCT predictable sequence task (Fig.3.b) that the NR population mean activity, black lines (Fig.3.b.2-6), become more and more linearly increasing with time to the next trial, which is a rewarding trial, thus the full motor cortical population average activity is tracking the time to the next R associated CS starting from the previous trial. This population activity peaks post R-CS, as seen in the OT-CCT R trials (Fig.3.b.2-6). This population tracking of the trial sequence is also clear in Fig.3.b.1 where we have plotted the % units in the full population that show significant differences between R and NR trials. Note, that many units show separability pre-CS in Fig.3.b.1 as compared to the manual task (Fig.3.c.1), which had a random trial value sequence. For both the manual and OT-CCT task there is an increase in the number of units that show significant differences between R and NR trials from session 1 to sessions 2, and onward, indicating the animal is learning the association between the CS and the reward (US, unconditioned stimulus) regardless of the environmental stability, that is fully predictable vs. random trial value sequences (Fig.3.b.1-c.1, sup fig.2). Both the manual and OT-CCT task data show two peaks in these % unit plots (Fig.3.b, c.1, sup fig.2), one post cue (CS) and one post reward delivery. Differences between the manual and OT plots are likely due to uncertainty, from the trial sequence, and also manual errors present in the manual task vs. the OT task. The results in Fig.3 were recorded from M1 contralateral to the arm used by the NHP in the manual CCT. We hypothesized that the reward modulation signal would be broadcast to M1 in both hemispheres and show support of this in sup fig.1-2. The results shown in Fig.3.b.1 indicate that the M1 population is clearly separable between R and NR trials even before the CS is shown due to the predictable sequence of the trial’s value, that is R trials followed by NR and repeating.

**Figure 3:**
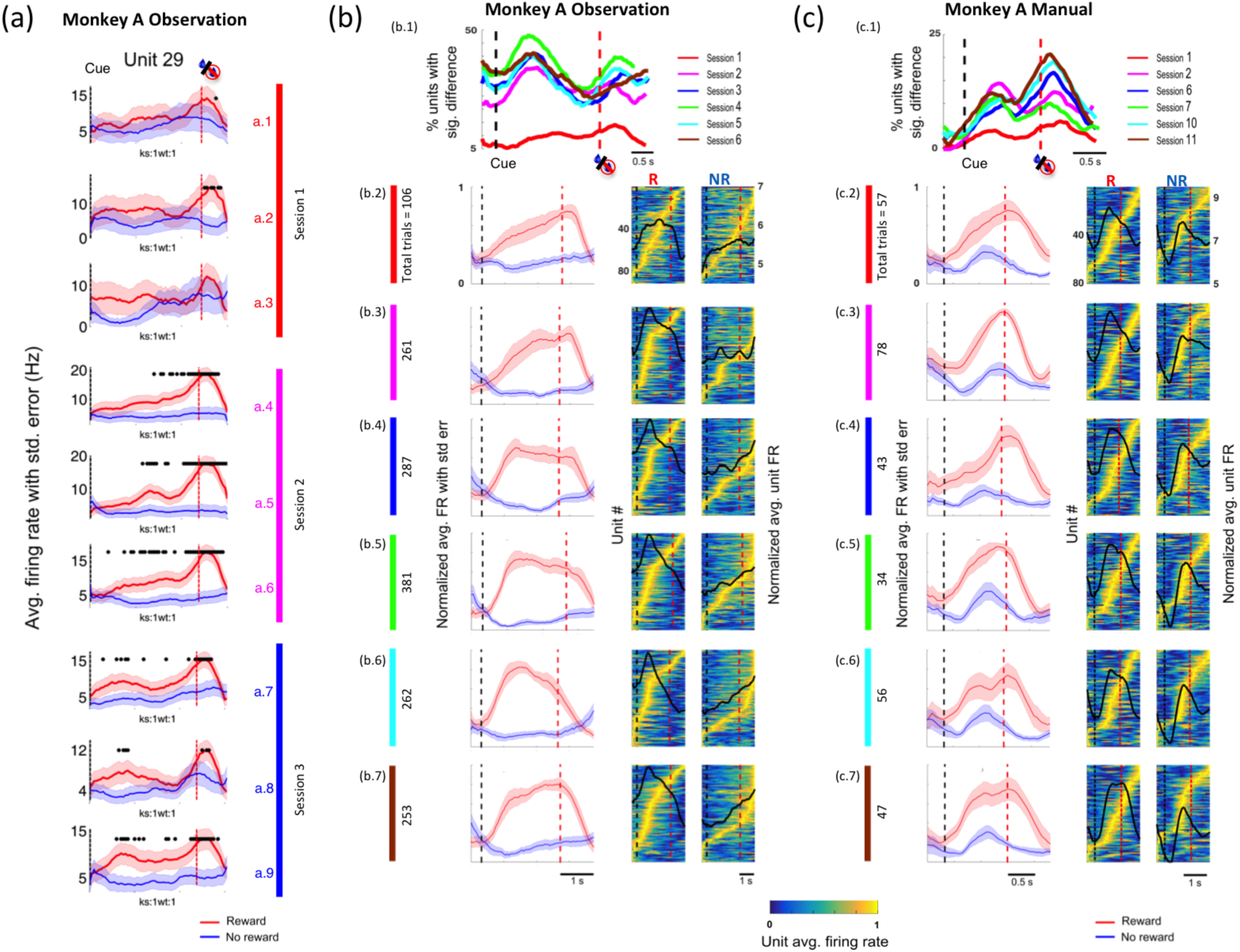
Single unit and population activity showing state value function evolution during learning. (a) Single unit example post-cue-time-histograms (PCTH) for rewarding (R, red) and non-rewarding (NR, blue) trials from the observational CCT task (OT-CCT) utilizing a fully predictable trial value sequence of R-NR repeating. Shown are PCTHs taken from three separate sessions (red, pink and blue vertical bars), with each session further divided into three sub-sessions. Unit 29 had a statistically significant difference in its median activity (Wilcoxon rank sum test, p<0.05) across R and NR trials earlier in the trial as we progressed to the later sub-sessions of the day. (b.1,c.1) Percentage of units that have significantly different (Wilcoxon rank sum test, p < 0.05) median firing rates between R and NR at a given time point (i.e. bin number, bin = 50ms). They also showed an increase in the percentage of units with significantly different median activity across R and NR trials post cue in session 2 and later compared to session 1. Therefore, the M1 neural ensemble represents a reward signal which is increasingly predictive of the yet to be delivered juice reward as we progress to the later sessions of the day. (b.1) OT-CCT data used for plots. a (c.1) Manual CCT utilizing a random trial value sequence. (b.2-7, c.2-7) First column; sub-population PCTH, red (R) and blue (NR) line plots. The data used for these plots was from the top 10% of units with respect to their correlation with the reward sequence (see *Methods*). These plots show that the peak of the normalized average firing rates measured across the top 10% units shifted from the reward (juice) delivery time point to right after the cue. The second and third columns include data from the full populations as PCTH in false color with black lines showing the mean of the full population. Binning of the neural data was performed at 50ms. Normalization of the average activity of a given unit was performed resulting in a range from 0 to 1 (see *Methods*). Units in the PCTHs in the second and third column were sorted for each session in decreasing order with respect to the time a given unit reaches the maximum average firing rate across rewarding or non-rewarding trials. These subplots show that the number of units that reach the maximum average firing rate across rewarding trials early in the trial increase as we progress from session to session on a single day.

### M1 learns and tracks the implicit value associated with various states in a task

In Fig.4 we present results from an uncued version of the GF task (Fig.1.b) with results consistent to that from the CCT task. The probability of being rewarded at the end of a successfully completed trial was dependent on the session trial value predictability (TVP). Chance trial value predictability (TVP) indicates no clear predictable trial value sequence structure, whereas complete trial value predictability was a simple alternating sequence of rewarding and non-rewarding trials. Therefore, TVP had to be inferred and tracked internally due to the absence of an explicit CS. In Fig.4, we present results from both the manual and observational versions of the uncued GF task. Few M1 units showed a significant difference in their activity across various time windows of the trial until the post reward window in sessions with chance TVP. In contrast, a large percentage of units had significant differences in their R vs. NR activity at the beginning of the uncued trials with complete TVP. This suggests that M1 inferred and tracked the underlying TVP in a given session and encoded the expected state value at the beginning of a trial based on the inferred trial value.

**Figure 4:**
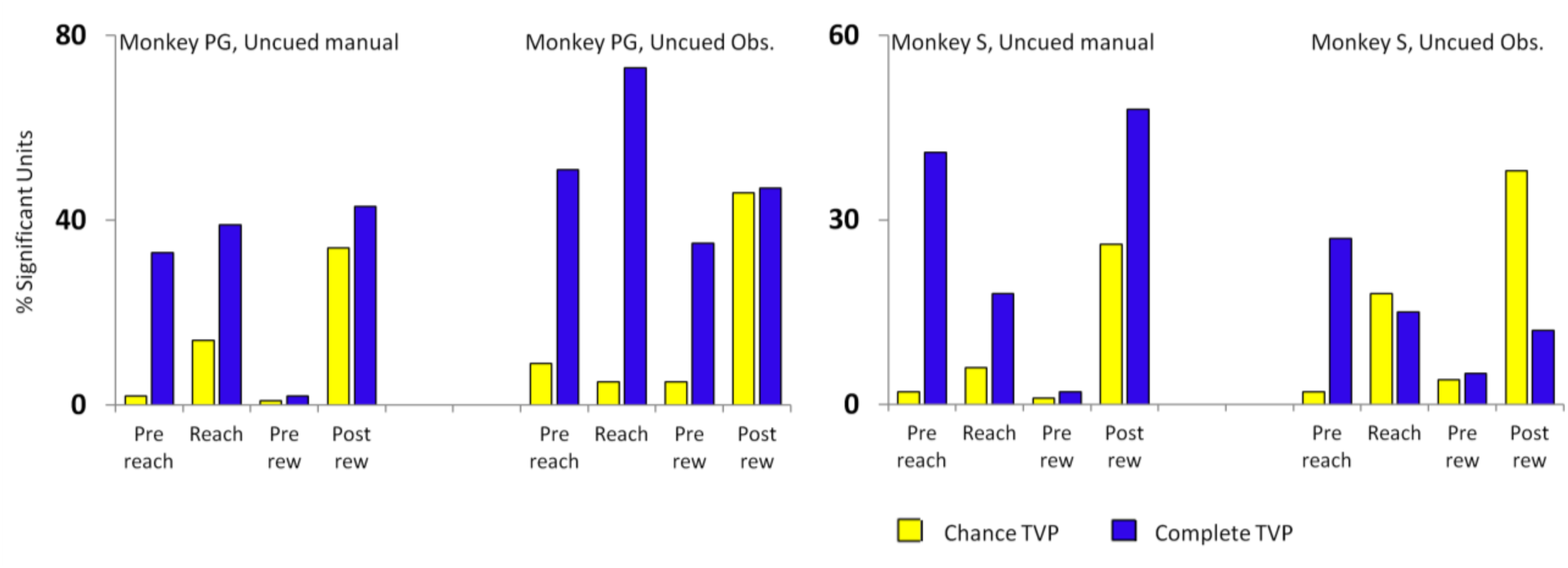
M1 tracks the state value without explicit conditioned stimuli. Two version of the uncued grip force task, manual and observational, with varying levels of trial value predictability are shown. NHPs conducted the grip force task seen in Fig.1.b without a conditioned stimulus that is without the green squares indicating the reward value of the trial. Shown for each NHP are results from the manual (left subplots) and observational (right subplots) versions of the task for NHPs PG (left two subplots) and NHP-S (right two subplots). Note that both NHPs show separability pre-reach and pre-reward for the complete TVP. When the trial values are unpredictable (chance TVP), and again there is no conditioned stimulus, i.e. uncued, the most separable trial time is post reward. Note that in the manual trials, even the complete TVP is highest post reward, as the NHPs do make errors and thus the value of the trial is only known with certainty post reward, while for observational trials the peaks are earlier in the trial before the reward.

### State value function of the MSTD model accurately captures reward related neural dynamics from M1

In order to further test our hypothesis that M1 holds a state value function we ran simulations of the microstimulus temporal difference (MSTD) RL model (20) using the same experimental structure as the real experiments (see Methods for model description). In Fig.5.a we have plotted the value functions from this MSTD model (Fig.5.a left column) as well as data from the NHPs Fig.3.b.2-4 for comparison. There is clearly a strong resemblance between the predicted value function by the model and the neural data (see sup fig.3-5 and the corresponding sections for further supporting results). The cross-correlation between the model value function and the neural data in Fig.4.a is on average r = 0.91. In Fig.5.b we have plotted the PCTH for example single units from M1 with an average cross-correlation between these units and the value function for the MSTD model of r = 0.92.

**Figure 5:**
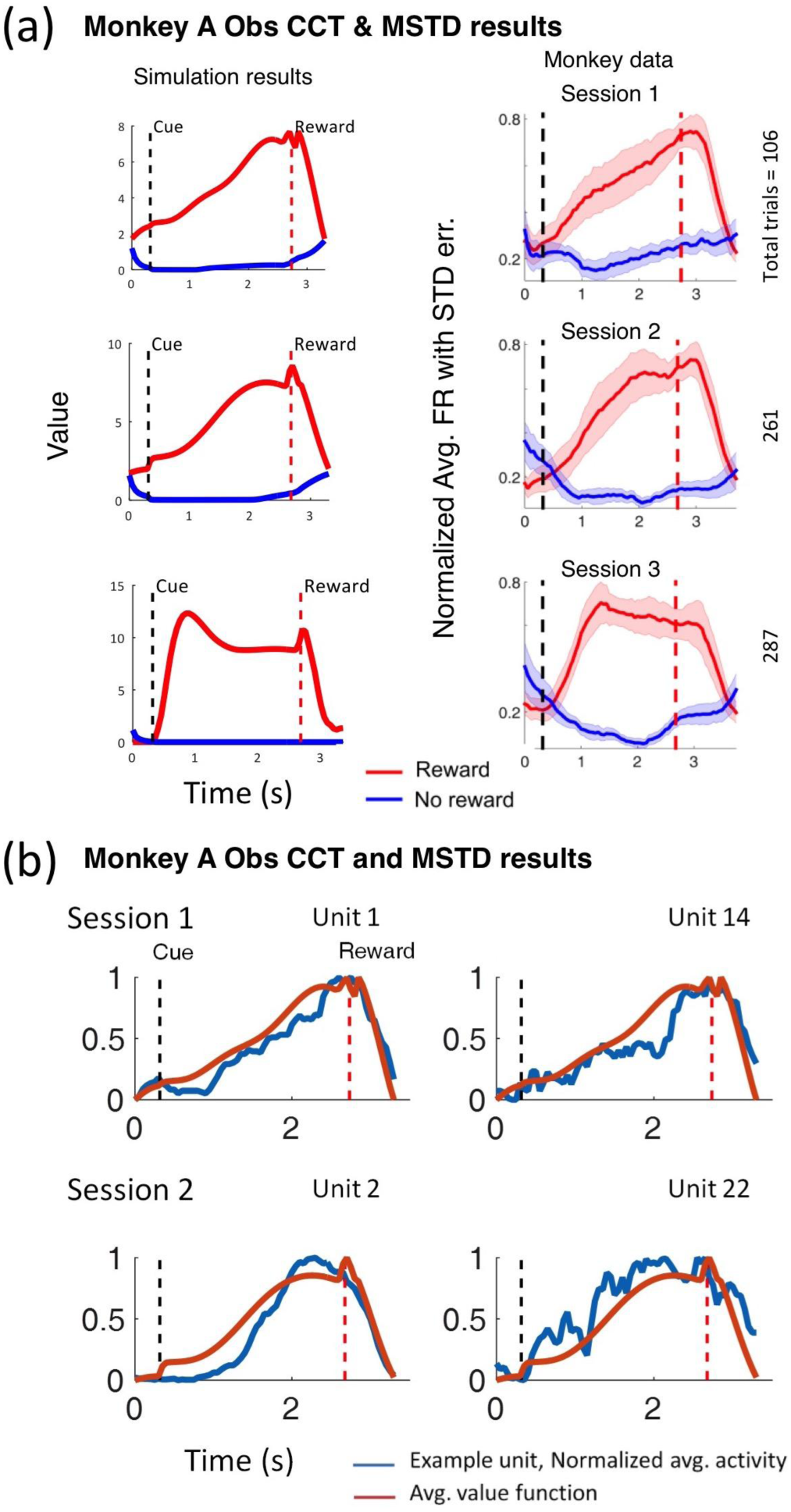
Development of the state value function during learning in a microstimulus temporal difference reinforcement learning model (MSTD) mirrors that of primate M1. (a) Left column subplots are the output value for the given trial time, x-axis, from mTDRL simulations run using the same state and reward sequences that the actual NHPs visited. The right column is the data from Fig.3.b.2-4 reproduced for comparison; briefly it is the normalized firing rate for a subpopulation from M1 while the NHP observed the task seen in Fig.1.a. (b) Single unit examples from M1 with the mTDRL value function overlaid for comparison.

### M1 encodes multiple levels of reward

NHPs S and PG performed the cued GF manual task with multiple levels of reward. A cue at the beginning of the trial informed the NHP of the amount of reward it would receive at the end of a successfully completed trial. The NHPs experienced approximately an equal number of trials for each reward level presented randomly. Here, we have plotted the average firing rate and standard error of the mean (window size ~ 1s post cue) for M1 example units with respect to the cued reward level. We found two types of units that linearly increased or decreased their firing rates with an increase in the amount of reward expected at the end of the trial (Fig.6). About two thirds of units that had significant fits with the reward value had positive slopes (linearly increased their activities with reward level) and the remaining one third had negative slopes (linearly decreased their activities with reward level). The M1 units encoded the expected reward post cue itself, i.e. before the reward was delivered in a given trial. The M1 ensemble encoded multiple levels of reward irrespective of whether the task was performed manually or passively (observed) by the NHPs.

**Figure 6:**
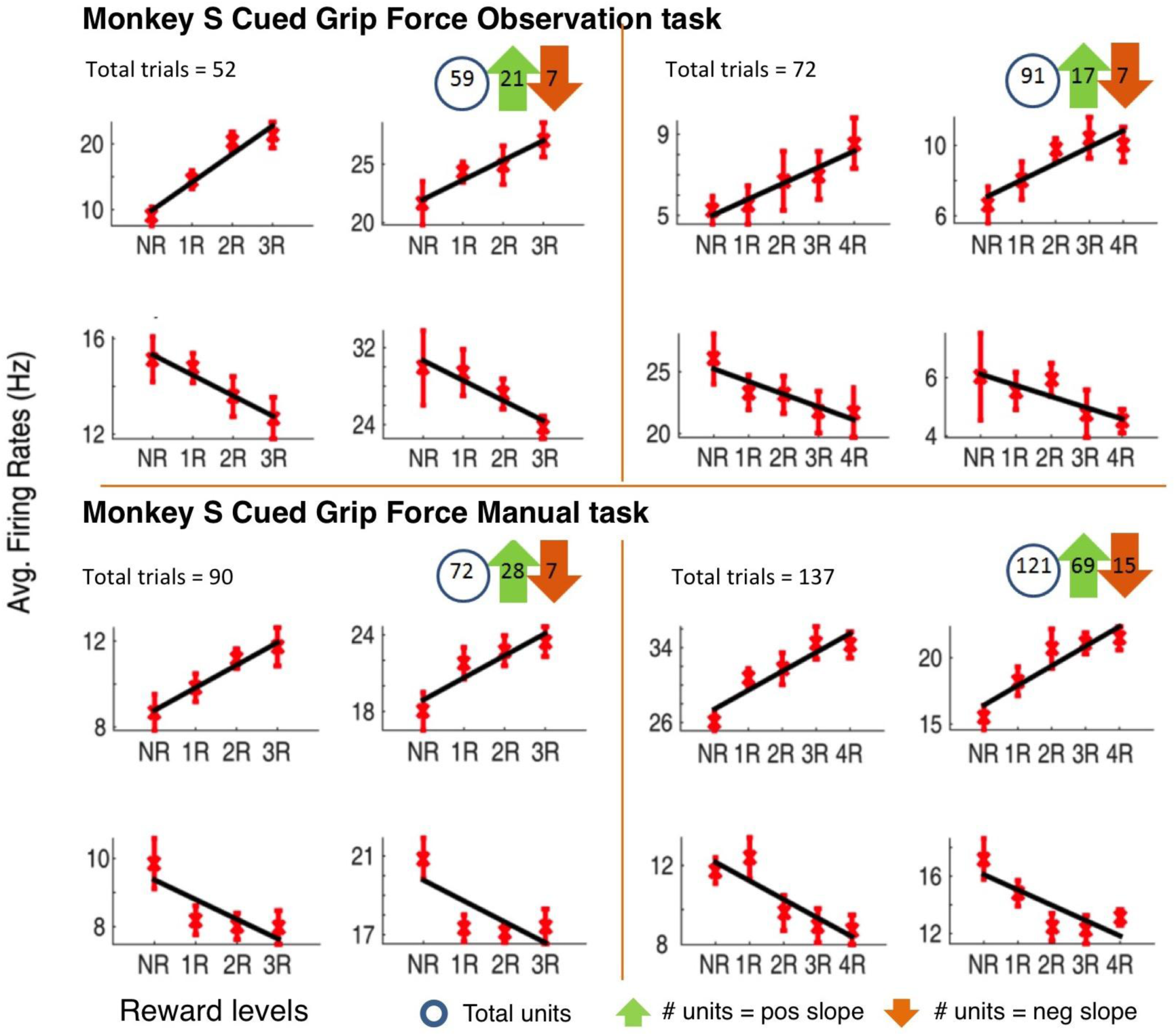
M1 encodes multiple levels of reward in a linear fashion. (a, b) Example M1 units’ mean firing rates, with SEM, for the cued reward value on the x-axis, from observational and manual GF tasks with multiple levels of reward for NHP S. The top row represents example units that increase their firing rates with respect to reward level, whereas the bottom row shows example units that decrease their firing rates with respect to reward level. The average firing rates were calculated from the time period of 500ms before reaching began to 500ms after the grasping scene in each trial. Approximately two thirds of units that had significant fits with reward levels had positive slopes (linearly increased their activities with reward level, the number of which indicated by green arrows) and the remaining one third had negative slopes (linearly decreased their activities with reward level, the number of which indicated by orange arrows). The circles represent the total number of units in the M1 ensemble.

### Reward prediction error: M1 units signal an omission of predicted reward

In order to determine if M1 has clear reward prediction error signals, we used reward catch trials in 10% of the trials in a set of sessions. In these catch trial sessions, a conditioned stimulus was used to signal rewarding trials as in our cued tasks, however, in 10% of the cued rewarding trials, we withheld the expected reward. The catch trials were dispersed randomly throughout the session. Such catch trials should introduce a discrepancy between the expected reward and the actual reward received. This discrepancy between the expected and actual reward is called the reward prediction error (RPE) and the TD form of this is considered one of the hallmarks of TDRL. Given that M1 neurons encode the expected reward value, one might also expect it to modulate its activity to an error in its expected reward. Ramakrishnan et. al. (15) have recently reported that the primary motor cortex encodes a RPE. Here we support the finding and extend it further to state that we see two unit subpopulations that modulated their activity in an opposite manner in response to a reward omission, with one population increasing activity and the other decreasing activity (Fig.7).

NHPs S and PG performed a cued GF task manually. The task consisted of reward (R = 0 or 1) and punishment cues (P = 0 or 1) resulting in one of four possible outcomes, depending on if the trial was completed successfully or unsuccessfully. A reward cue indicated that the trial would be rewarded if completed successfully, and the lack of a reward cue indicated that the trial would not be rewarded if completed successfully. A punishment cue indicated that punishment in the form of a five second timeout period would be delivered if completed unsuccessfully, and the lack of a punishment cue indicated that the trial would not be punished if completed unsuccessfully. These cues were presented at the beginning of the trial, informing the NHPs of the outcome contingency in that trial. Ten percent of trials were catch trials, where the expected reward based on the cue was not delivered. In this analysis, we considered regular trials as those cued rewarding, completed successfully, and with reward delivered, and catch trials as those cued rewarding, completed successfully, but with no reward delivered. M1 units were sensitive to the omission of the predicted reward (Fig.7). There were two types of units, those that significantly elevated their activity (Fig.7.a and first column of Fig.7.b) and those that significantly depressed their activity in response to a reward omission (second column of Fig.7.b). The percentage of units from the total population that significantly increased or decreased their activity in response to a reward omission were 9% and 17% in NHP PG (total units = 82) and 81% and 1% in NHP S (total units = 79), respectively. Two subpopulations of units similar to what we have reported here in M1 were also reported in posterior cingulate cortex (CGp) by McCoy et. al. (21).

**Figure 7:**
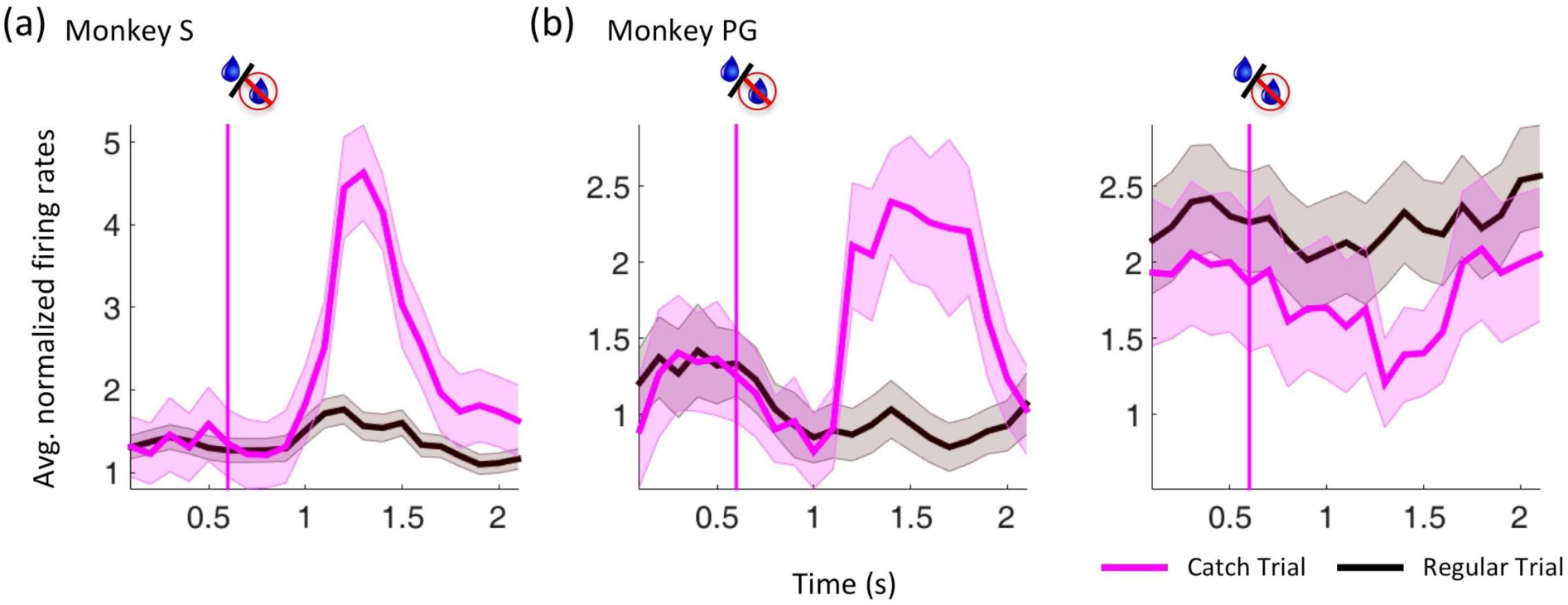
Reward prediction error. Two distinct populations of reward prediction error in M1. (a) and (b) show the PSTHs (mean and standard error) of the M1 subpopulation aligned to the reward delivery/omission for NHPs S and PG in regular and catch trials. The pink vertical line displays the usual time of reward delivery in R trials. The average activity across M1 subpopulation between regular and catch trials was significantly different (ttest2, p<0.05). Binning of the data was done at 100ms. (a) Average activity of M1 units that significantly increase their firing rate in response to reward omission. (ttest2, p<0.05) (b) Average activity of M1 units that significantly increase (1st col) and decrease (2nd col) their firing rate in response to reward omission. (ttest2, p<0.05)

## Discussion

In the work presented here we show in a variety of tasks that the primary motor cortex (M1) encodes a state value function that captures the state dependent temporally discounted expected reward, in line with that from TDRL. We show the temporal evolution of this state value function during the learning process of either an explicit conditioned stimulus that fully predicts the trial value, or during learning of predictable sequences of trial values. In addition we presented results showing there are two types of response in M1 to reward prediction error, both increases and decreases in firing rate. These results, which in general agree with our MSTD simulations, were consistent across 5 NHPs (4 trained, 1 naive to the tasks), while performing either a reaching task or a grip force task, and were observed bilaterally, irrespective of whether the tasks were performed manually or were passively observed by the NHPs. It is interesting to note that the MSTD model, which has been shown to accurately model dopaminergic neural activity in the form of a TD error, was also capable of accurately modeling, in the form of the state value function, the reward related activity observed in M1. Thus, M1 activity holds all the hallmarks of a temporal difference reinforcement learning model.

Previously our group has shown that reward modulates both the primary motor (M1) and somatosensory (S1) cortices during action and action observation (2,17,22), and that individual M1 neurons multiplex information on kinematics and reward (2). This work was subsequently supported by Ramakrishnan et al. 2017 (15) where they also showed that M1 and S1 responded to reward prediction errors and that reward could modulate sensorimotor directional tuning curves. We recently showed that this reward modulation of directional tuning holds during intracortical brain computer interface control (iBCI) when utilizing M1 (23), and that M1 grip force tuning functions are modulated by reward during manual control (23). In addition, we have recently discovered that reward modulation influences the relationship in M1 between the local field and individual neurons, where the spike field coherence is upregulated during non-rewarding trials as is the phase amplitude coupling between alpha phase and gamma amplitude (24).

In reinforcement learning, reward prediction error (RPE) is the difference between expected and actual received reward, where these expectations are held by the state value function. Recognizing such an error and utilizing it to modify the state value function drives learning, and state values drive action selection, or behavior. Such learning helps animals better predict their possibly non-stationary environment. Delivery of an actual reward that is better than expected results in a positive RPE, and similarly a negative RPE indicates that an outcome was worse than expected. Positive RPE tends to result in increased activity in DA neurons whereas negative RPE tends to result in a depression of DA activity (9,15,22). In our uncued work, the absence of a reward predicting cue (CS) and chance TVP led to a situation where the NHPs were incapable of perfectly predicting whether the current trial would lead to reward or not. Thus, they could not build up a strong state value function. The M1 response observed at reward delivery time in such scenarios may encompass a RPE, and a response to the primary reward as well. We continued to observe elevated M1 activity to reward delivery even after extended exposure to reward, or in experiments where reward delivery is predictable either due to a higher than chance TVP and/or due to the presence of a CS. Therefore, it is unclear at this point whether the M1 response observed post/during reward delivery was solely representative of the value of the reward delivered, about to be delivered, or is also encoding RPE. Past these influences on the neural activity there is little doubt that attention and motivation are also influencing the neural population and are modulated by the reward landscape. These concepts have been treated previously and our future work will more directly address these variables (18, 25-26). It is difficult to see how motivation alone, or attention could explain all of our results. However, as motivation may also be carried by part of the dopamine signal (13), at least part of the reward modulation of M1 is likely due to this dopaminergic motivation signal, which in turn may modulate attention, as would the value function one would expect. Additionally, serotonergic neurons from the dorsal raphe nucleus have been shown to encode reward (27,28). How these signals interact with dopaminergic reward signals, and if they might encode something broader such as the “beneficialness” of a particular environmental context remains to be seen (29,30). Importantly though, a subpopulation of serotonin neurons in this region has been shown to increase firing rate in response to a cue, and then further increase firing rate in response to reward delivery in a fully learned task (27). This, and the fact that similarly to midbrain dopamine these neurons project widely throughout the cortex (31), suggests that serotonin may be the cause of some of the modulation seen here, particularly in the post-reward delivery period.

A key component of RL is the state value function that captures the expected temporally discounted reward, from a given state in the environment. The value function can also be modified by the animal’s knowledge and certainty of its environment. The value function, unlike TD error, is a slower varying function in nature existing throughout the trial, similar to the reward signal we have presented from M1. The value function under the MSTD model shows that the RL model is capable of predicting the expected reward even in the absence of a conditioned stimulus as long as the reward predictability was higher than chance. The value function under the MSTD model also evolved across trials in a fashion similar to that observed in the M1 reward signal across multiple sessions in the NHPs (Fig.5). Similar behavior of the state value function was observed bilaterally in the NHPs’ M1, (NHP A – contralateral to the right hand, NHP Z – Ipsilateral to the right hand) even though the NHPs always performed manual tasks with their right hand. The value function under the MSTD model also became increasingly predictive of the expected reward even before the presentation of the cue with increased reward predictability. Such behavior was also observed in the M1 ensemble activity. One can imagine that the NHPs must have an internal representation of the value of the juice reward being awarded at the end of a successfully completed trial. This is observed even more clearly in sessions with multiple levels of reward. The M1 ensemble represented the expected reward right after the presentation of the cue thus suggesting that the motor cortex isn’t only encoding whether a reward is being awarded or not but also capturing the expected reward level. The value function under MSTD model is also capable of capturing multiple levels of reward. Therefore, we propose that the reward signal in M1 is reminiscent of the value function learned under the Microstimulus Temporal Difference (MSTD) Reinforcement Learning model, and by extension suggest that the primary motor cortex encodes a value function.

The work presented here has applications towards autonomously updating brain machine interfaces (32-36). Although the idea of replacing the lost or damaged limb with a prosthetic device is not a new concept, only recently has the technology been developed to allow the control of these devices via a neural signal from the user. Sensorimotor brain machine interfaces strive to integrate the sensorimotor system and a neuroprosthetic thus providing people with sensory (37-43) or motor disabilities the capacity to interact with the world hopefully in a more complete manner. Various sensorimotor BMI algorithms and architectures have allowed animals and humans to control external devices (44-50), where in these systems sensory feedback was via the intact visual system. Typically BMI systems utilize an exact error signal to adapt the BMI using supervised learning. This generally requires a controlled environment such as a laboratory setting, and thus may restrict the usefulness of these methods in the complex, evolving environments that we live in. Additionally, neural input to BMIs change with learning and time due to inherent instabilities such as the loss of single units or addition of new units, as well as due to the learning of state values as described in this report. It is also known that such value encoding changes M1 representations of directional tuning and grip-force turning functions (23). The concept of reinforcement learning may prove to be useful in transitioning BMIs to novel and unstable environments (32-36), as RL doesn’t require a full error signal such as that used for supervised learning to update BMIs. Particularly useful may be the actor-critic reinforcement learning architecture (6), where the actor is the portion of the system that decodes the instructional neural activity into actions and the critic decodes the evaluative neural signal critiquing the action performed to update the actor. One can put these ideas in terms of making a reaching movement. If the actor decodes a time bin of instructional neural activity as indicating a rightward movement, which the BMI agent then executes, but the evaluative neural signal after this movement indicates “things” are not going well, then the BMI system can learn that such instructional neural activity should not be interpreted as moving to the right in the future. Likewise, if the actor made the correct move, which would be seen in the evaluative neural feedback, this could be used to increase the likelihood of making that movement when the corresponding instructional neural pattern is encountered in the future. In this manner the system will automatically update itself when there are changes in the neural input to the system. Our current and future work is towards developing BMIs that work on these RL principles and utilize the state value information to stabilize the BMI against changes in value/motivational based neural directional tuning.

## Methods

### Cued-Center Out-Reaching task (CCT)

NHPs sat in a primate chair with their right (dominant) arm in an exoskeletal robotic manipulandum (KINARM BKIN). NHPs A (male, *Bonnet Macaque*) and Z (female, *Bonnet Macaque*) were proficient in performing an 8-target center-out reaching task before implantation. NHPs A and Z were implanted in the contralateral and ipsilateral M1 (with respect to the right arm) respectively. These NHPs were then introduced to the cued center out reaching task (CCT) (Fig.1.a) (2). A hand-feedback cursor was displayed on a screen in the horizontal plane just above their right arm in alignment with their right hand during the manual task. The NHPs performed cued reaching tasks, where the reward level was cued via the color of the reaching target. Progression from the current trial to the next was only allowed following successful completion of the current trial. In an observational version of the task the NHP passively observed constant speed cursor trajectories to the cued target. The data considered here from NHP A corresponds to the days when it was relatively new to the CCT, on the sixth day of manual and observation CCT. NHP A performed manual and observational CCT with chance and complete trial value predictability (TVP). The chance TVP session had a random sequence of R and NR trials in a given session, whereas in the complete TVP session a R-NR sequence was repeated in a completely predictable manner. The data considered in this paper from NHP Z are from days with a bias in the number of R to NR trials presented randomly (66% R trials). These sessions were from days after NHP Z had a break of 20 days from performing the CCT, during which it performed brain machine interface tasks. In addition, NHP Z did multiple types of tasks on these days, including CCT, manual COR, and BMI experiments. NHP B, our pseudo-naive male NHP (*Bonnet Macaque*), was unsuccessful in learning the manual COR task. He did not perform a single successful manual reach to the target in 18 days of training spread across 2 months. We stopped the training at this point and considered it a naive animal, which is supported by the near zero R values for predicted kinematics from its neural data. The data considered in this work from B corresponds to the second day ever of it experiencing the observational CCT. The NHP was required to maintain its gaze on the task plane throughout the trial period via eye tracking, as the virtual cursor moved only when gaze was maintained on the task plane.

### Grip Force (GF) task

NHPs S (male, *Bonnet Macaque*) and PG (female, *Rhesus Macaque*) were required to apply and maintain an instructed (with tolerance) amount of grip force (applied force shown as red bars, instructed force shown as blue bars Fig.1.b), from initialization of the grasping phase until the robot automatically moved the cylinder to a target position. NHPs were required to release the grip to successfully complete the task and receive or not receive juice based on whether the trial was rewarding or non-rewarding, respectively. Progression to the next trial required successful completion of the current trial. The difference between cued and uncued GF tasks was the presence of a conditioned stimulus (CS) indicating the trial’s value. The presence or absence of a cue at the beginning of a cued GF trial informed the NHP on whether it would be rewarded or not at the end of a successful trial. There was no CS during uncued GF tasks. Both NHPs were required to perform the grip manually in the manual version of the task, and they passively observed while the grasping was performed automatically in the observation version of the GF task. All NHPs performed the grip force task with their right (dominant) hand. The GF task had the following 6 stages. 1.) Cue - This stage is observed only during the cued GF task. A cue (CS) explicitly informed the NHP of the reward value it would receive at the end of a successfully completed trial. Cues flew into position at the top of the task plane during this stage. The cue was maintained in the task plane throughout the trial period; no cue flying in equaled a no reward task. 2.) Reaching - The cue presentation was complete before the task entered into this stage. The virtual robot moved automatically at a constant speed from the rest position to the cylinder location. 3.) Grasping – The NHP applied grip force, virtually represented in real time as a force bar in red, its width proportional to the amount of force exerted on the physical manipulandum (Fig.1.b). The blue bars in Fig.1.b represent the instructed force to be applied and maintained during the task by the NHP. The width of the blue bars instructed the tolerance allowed in matching the applied grip force to the instructed force. The grip force was considered valid as long as the applied force (red bar) was within the lower and upper boundary of the blue bars. Over or under application of force resulted in failure and a repeat of the same trial value. During the observation GF trials the “grip force” is applied automatically, and the NHP was required to passively watch the task being performed. 4.) Transport – The virtual robot automatically moved the cylinder at a constant speed from the start to the target position, given that the NHP maintained the instructed force. 5.) Release – The NHP was required to release its grip to successfully complete the trial and either receive or not receive juice reward. The release scene was automatically executed during the observational GF trials. 6.) Reward-Juice was delivered to the NHP following a successfully completed rewarding trial, and no juice was awarded following successful non-rewarding or unsuccessful trials.

### Surgery

Electrode array implantation was performed after the NHPs manually performed the GF tasks (NHPs PG and S) or the COR tasks (NHPs A and Z) proficiently as described in the earlier sections. NHP B was implanted having never successfully performed a COR task. Chhatbar et. al. (51) describe the implantation procedure in further details. Electrode arrays were implanted in M1 either contralateral (NHPs S, PG, A and B), or ipsilateral (NHP Z), to the right arm used by the NHPs to perform the CCT or GF tasks. In short, the array implantation procedure was as follows - 1) dissection of the skin above the skull 2) craniotomy 3) durotomy 4) cortical probing in the primary sensory cortex (S1) to accurately locate the hand and the arm region 5) implantation of the electrode array in the forearm and hand region of the primary motor cortex and 6) closure. Following craniotomy and durotomoy, markers such as the Central Sulcus, Arcuate Sulcus, Arcuate Spur and the Intraparietal Sulcus were used to confirm our location on the cortex. Cortical probing was performed using Neuronexus silicon neural probes. They were inserted into S1 and real time neural activity was heard on speakers while a surgery assistant touched the right/left arm and hand of the NHP. Once the hand and the arm regions were recognized in S1, a chronic 96 channel platinum microelectrode array (Utah array with ICS-96 connectors, 1.5mm electrode length, Blackrock Microsystems) were implanted in the primary motor cortex mirroring the S1 forearm and hand region across the Central Sulcus. All surgical procedures were performed under the guidance of the State University of New York Downstate Medical center Division of Comparative Medicine (DCM) and were approved by the Institutional Animal Care and Use Committee in compliance with NIH guidelines for the care and use of laboratory animals. Aseptic conditions were maintained throughout the surgery. Ketamine was used to induce anesthesia, and isofluorane and fentanyl were used to maintain the animal under anesthesia during the surgery by and under the guidance of DCM. Possible cerebral swelling was controlled by the use of mannitol and furosemide whereas dexamethasone was used to prevent inflammation during the surgery. A titanium post (Crist Instruments) was implanted on the skull with an attached platform built in-house to mount the 4 ICS-96 connectors from the Utah arrays (51). Antibiotics (Baytril and Bicilin) and analgesics (Buprenephrine and Rimadyl) were administered in line with the DCM veterinarian staff recommendations.

### Electrophysiology

NHPs were allowed to recover from surgery for 2-4 weeks. Following the recovery period, single-and multi-unit activity and LFPs were recorded from M1 while the NHPs performed the behavioral tasks. Multichannel Acquisition Processor systems (Plexon Inc.) were used to record neural data. The unit activity was captured at a sampling frequency of 40KHz. Offline sorting of the neural data was performed using template sorting in Offline Sorter (Plexon Inc.) to identify individual single-and multi-units. Parameters of the templates used for sorting in the first file of a day were saved and used for sorting remaining data from that day. This provided us with units across multiple files on the same day that had similar sorting templates and parameters.

### Microstimulus Temporal Difference model (MSTD)

The reinforcement learning problem, for optimal control, is for the agent to maximize its cumulative temporally discounted reward from the environment. The agent uses information, such as from sensory systems, which we call states, in order to complete this reward optimization task, that is if the agent is actively choosing actions, such as in our operant conditioning tasks (manual tasks). During our observational tasks (classical conditioning) the agent can still use the state information to build state/value associations that is the state value function. Dopaminergic centers of the brain have been shown to represent reward probabilities, value of reward predicting stimuli and error in reward expectation (1,14,52). Recent work has reported a ramping up of dopaminergic activity as an animal approached its goal i.e. reward (53). All such modulations observed in the dopaminergic centers have been modeled and predicted well using basic and modified Temporal Difference (TD) reinforcement learning models (20, 54-56).

In most trial and error learning scenarios rewards are delayed with respect to the actions that caused them, or the states that predict them. This leads to what is known as the credit assignment problem, which describes how an agent knows what actions and states to assign the credit for later rewards. Under the basic TD model, the stimulus, such as the CS in our tasks, is represented as a complete serial compound (56), suggesting that the agent is aware of the exact amount of time that has elapsed since the CS. Such an assumption led to an incomplete encapsulation of dopaminergic neurodynamics, especially when the reward timing was varied (20). Therefore, the assumption of a perfect clock in the basic TD model was replaced with a coarse temporal stimulus representation captured using a temporal basis set representation in the microstimulus TD model (MSTD) (20,57). The MSTD model utilizes a coarse temporal stimulus representation to overcome the shortcomings of the basic TD model. The MSTD model suggested by Ludvig et. al. (20) was tested and expanded in our current work, using simulations which mimicked various experiments performed by NHPs A, Z, B, S and PG.

Under the MSTD model, a stimulus onset results in a decaying memory trace for the corresponding stimulus. Such a trace when encoded with equally spaced temporal basis functions results in what were termed microstimuli. Each microstimulus’ value at a given time point keeps track of how close the memory trace height of a given stimulus is to the center of the corresponding basis function. Therefore, it acts as a temporal proximity measure and as a confidence measure that the memory trace has reached a certain height, or in other words, that the stimulus happened a given period of time ago. For further details please refer to (20).

The basis functions are defined as Gaussian functions

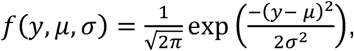

where *y* is the trace height, *μ* is the mean, and *σ* is the standard deviation of the particular Gaussian basis function. Each stimulus (CS for R (CSR) and NR (CSNR) in cued task simulations, US for R (USR) and NR (USNR) in cued and uncued task simulations) has its own memory trace and associated microstimuli. The trace height *y* of the memory trace was set to 1 at the onset of the corresponding stimulus and decayed at a rate of 0.985 on each time step following the stimulus onset. The level of the i^th^ microstimulus for a j^th^ stimulus at time t is given by:

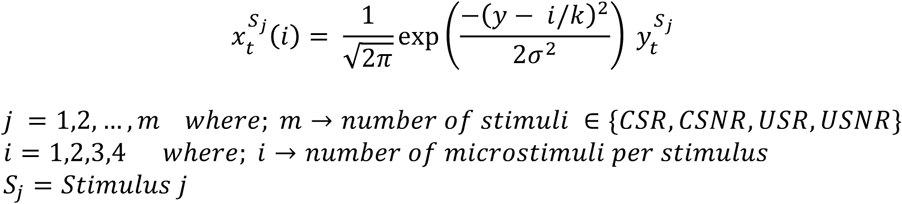

The coarse representation of the trace height, indirectly capturing the time since the stimulus, was then calculated using the above equation. The model attempts to learn the optimal estimate of the current state value. The state value can be thought of as a summation of the discounted future reward given the current time step in the trial, assuming the agent discounts reward value with the passage of time spent waiting for the reward. Given the microstimuli levels, state values were estimated as an absolute value of the linear combination of weighted microstimuli levels. The vector of adaptable weights mapping the microstimuli levels to the state value estimates are represented as *W*. The state value estimate at a given time step is:

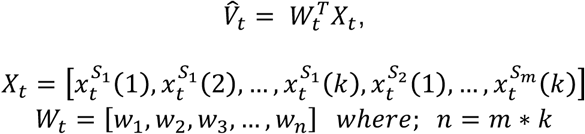

where n is the total number of microstimuli across all stimuli. Our simulations for cued experiments had 4 stimuli (CSR, CSNR, USR and USNR) with 4 microstimuli for each stimulus, resulting in m = 4, k = 4, and n= 16. Similarly, our simulations for uncued experiments had 2 stimuli (USR and USNR) with k = 4 for each stimulus resulting in n = 8, i.e. 8 temporal basis functions. The lower boundary for the state value is maintained at 0 resulting in no negative state value.

The TD error, which encapsulates the error in the estimated value, at every time step is calculated as follows:

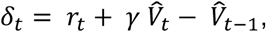

where *r*_*t*_ is the actual reward awarded during the reward outcome period of the trial, 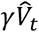 is the current discounted estimated value, and 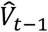 is the previous estimated value. TD error is then utilized to update the weights, which map the microstimulus values to the estimated state value 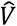 as shown below.

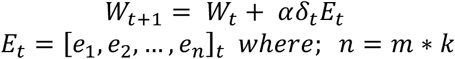

*α* is the learning rate and e are the eligibility traces (6) updated as follows:

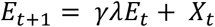

Eligibility traces are necessary for faster learning in a temporal credit assignment problem. These allow the propagation of the sparse rewards in the environment to the rest of the experienced state space. The decay of the credit such that the recently visited states are assigned more credit is represented by a decay factor *λ*. *γ* is the discount factor, encoding how fast the future rewarding events lose their value with time. The MSTD model was validated as detailed in the supplementary MSTD model validation section.

### Correlation of units with ‘reward’

A variable ‘reward’ was defined such that +1 or -1 was assigned to each bin of R and NR trials respectively. Data from cue presentation to the corresponding trial completion was considered for each trial. The correlation coefficient (*corrcoef*, MATLAB) was computed between each unit’s firing rate and the variable ‘reward’ in a given session. The unit with the highest positive correlation coefficient was most correlated with rewarding trials whereas the unit with the highest negative correlation coefficient was most correlated with non-rewarding trials.

### Normalization

Neural data acquired while NHPs performed various experiments were binned at 50ms. The average activity across R and NR trials for each unit was casually smoothed (window of 500ms) and concatenated to form a vector *Y* for a given unit. Subsequently, normalization was performed using equation shown below, resulting in a range from 0 to 1. ‘*Normalized y*’ was decatenated to obtain the normalized average activity of the same unit across R and NR trials. The process was repeated for all units in the neural ensemble.

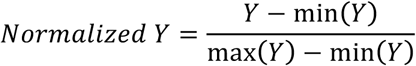

### Peri-cue-time-histogram (PCTH) of all units with false colors

Binning of the neural data was performed. The casually smoothed average activities across R and NR trials for each unit were normalized individually. Units were sorted for each session in decreasing order with respect to the time required for a given unit to reach the maximum average firing rate across rewarding or non-rewarding trials. Therefore, units at the top of the PCTH ‘neurogram’ reach the maximum average firing rate across trials later in the trial, whereas the units at the bottom of the neurogram reach the maximum average firing rate earlier in the trial. Units for R trials were sorted such that units at the top reached the maximum average firing rate across R trials later in the trial and similar logic was applied to the PCTH for NR trials. There was no requirement for maintaining the sorting order across sessions.

## Acknowledgement

We would like to thank Dr. Elliot Ludvig for his invaluable insights and discussions on the paper and the microstimulus temporal difference model (MSTD).

Research was supported by NIH 1R01NS092894-01, NSF IIS-1527558, DARPA REPAIR Project N66001-10-C-2008, NYS SCIRB contracts C30600GG, C030838GG, and DOH01-C32250GG-3450000.

## Supplementary Materials

Here in this supplementary material we present further support that M1 activity presents all the hallmarks of a temporal difference reinforcement (TDRL) learning system. We present further results from the NHPs and from the microstimulus TDRL (MSTD) model that corroborate the NHP data from a multitude of tasks, in both hemispheres, and for a task naive animal.

### The value function in M1 is represented bilaterally and in a naive NHP

NHP Z was proficient in performing an 8-target center-out reaching (COR) task before implantation, and was implanted in M1 ipsilateral to the right arm. NHP Z also had some experience with the cued center out task (CCT) post implantation. The results shown here for NHP Z are from sessions with an inherent bias in the number of R trials to NR trials in a session (66% R trials). In a given recording day, NHP Z performed brain machine interface (BMI) tasks, manual tasks, and the CCT. We show here that the reward related dynamics in NHP Z’s ipsilateral M1, performing the CCT, followed a similar trend as observed in NHP A (main paper, Fig.3) who was implanted contralaterally. For NHP Z the data sessions come from 3 separate days due to the fact that Z was also part of BMI experiments on the same days as the CCT recordings. (see Supp.fig.1 below)

NHP B underwent training to perform a center out reaching (COR) task manually for 18 days across two months. NHP B never performed a successful manual COR trial by itself during the training period. NHP B was then implanted in the contralateral M1 approximately two months post the cessation of training. After a four-week recovery period from the surgery, it then performed the observational CCT tasks. Therefore, we consider NHP B to be a pseudo-naïve animal. We investigated whether a reward signal would be observed in NHP B and if its M1 also represented reward related dynamics in a similar manner to NHPs A and Z, who were trained to manually perform the COR task. We show results from the second day of NHP B performing the observational CCT task, which was the first day where it was required to maintain its gaze on the task throughout the trial, from initiation of cursor movement post color cue till the time the virtual cursor reached the target. The cursor was allowed to move only when the NHP maintained its gaze on the task plane. This was done to ensure that the cues and cursor movements were in the visual field of the NHP and by extension that the animal was attending to the task.

In Supp.fig.1 we have plotted the average neural activity for a subpopulation of single and multi units that correlated with reward value, where R and NR trials are indicated in red and blue respectively (Supp.fig.1.a-b.1-3). Specifically, this subpopulation is the average of the top 10% of the population after rank ordering with respect to individual unit’s correlation with reward as was presented in the main text Fig.3. In both the manual (Supp.fig.1.a) and Observation (OT) (Supp.fig.1.b) versions of the CCT task, we see that the differences between R and NR trials are initially around the time of reward delivery, and this difference grows earlier in the trial over time as the subject continues performing the task. We have plotted the peri-cue-time-histogram (PCTH) for R and NR trials for all units as false color plots in Supp.fig.1.a-b, c.1-3. In each false color subplot, the black line is the mean of the population. Learning also took place for the population mean with increased trial experience. An increase in the percentage of units with significantly different activity at each bin between R and NR trials was observed earlier in the trial on day 2 and 3 compared to day 1 (Supp.fig.2). Therefore, the value function in M1 is represented bilaterally and in an untrained naive NHP.

**Figure 1:**
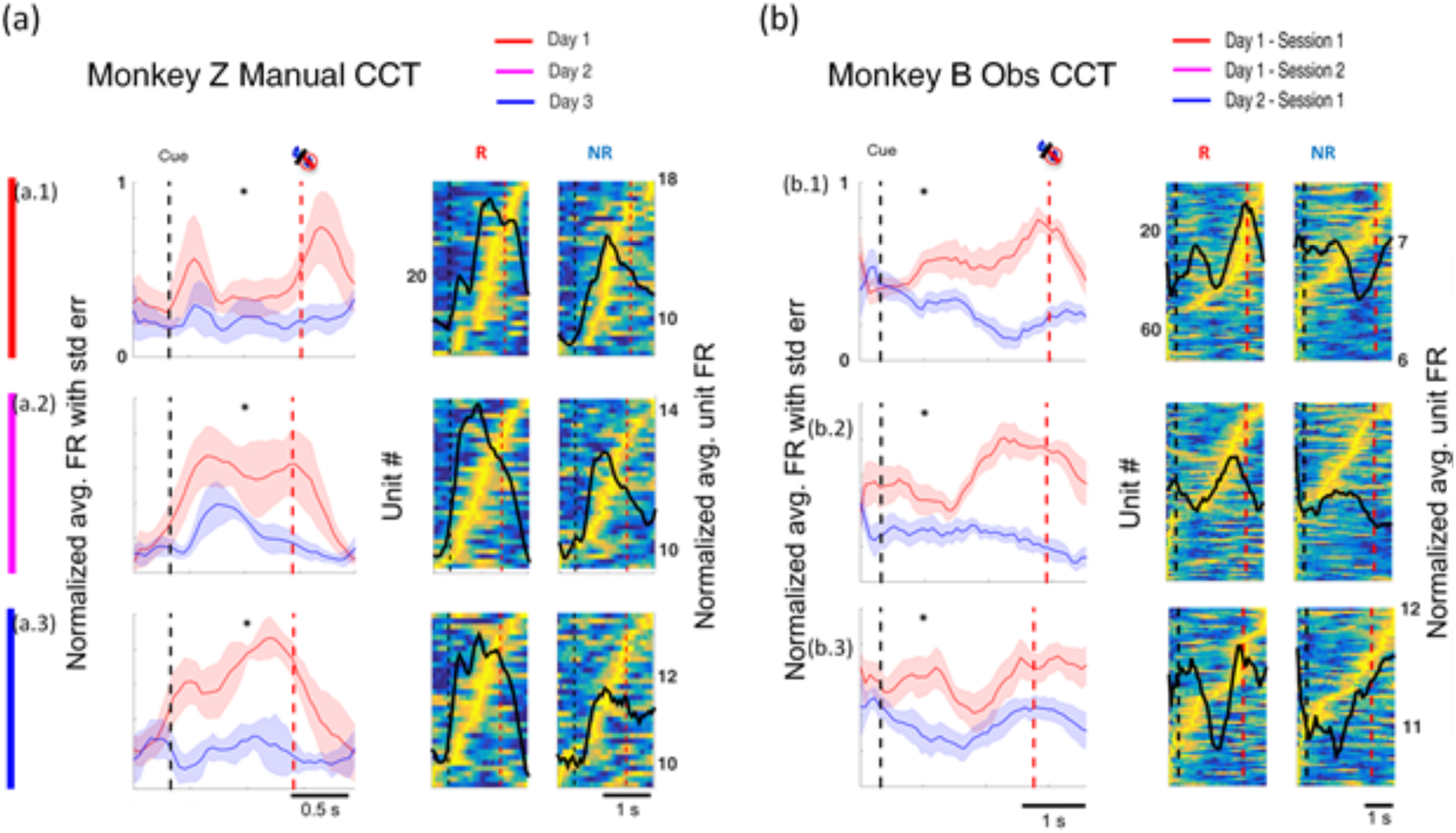
Population activity showing value function evolution with learning in NHPs Z and B. In the cued center out reaching task, NHP Z performed with the right arm, and NHPs Z and B were implanted in the primary motor cortex ipsilateral and contralateral to the right arm, respectively. Subplots a.2-4, b.2-4 depict the subpopulation PCTH of the top 10% correlating units, with rewarding (red) and nonrewarding (blue) line plots, and the full population PCTH in false color with the black line showing the mean of the population.

**Figure 2:**
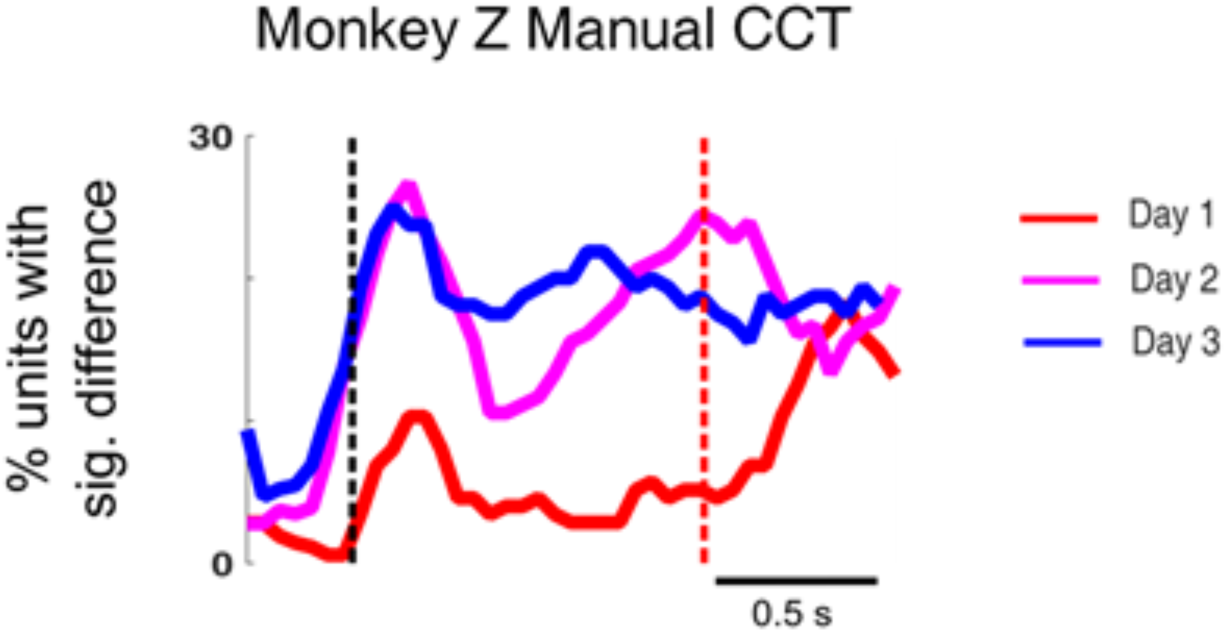
Percentage of units that have significantly different (Wilcoxon rank sum test, p < 0.05) median firing rates between R and NR at a given time point. This figure shows an increase in the percentage of units whose median activity across R and NR trials is significantly different (Wilcoxon rank sum test, p < 0.05) post cue compared to recording day 1. Therefore, the M1 neural ensemble represents a reward signal which is increasingly predictive of the to-be-delivered juice reward as NHP Z progresses from day to day.

### Microstimulus Temporal Difference (MSTD) value function predicts the expected reward value in both un-cued (no-CS) and cued (CS) tasks in NHPs

We performed simulations of the un-cued reward tasks with chance or complete trial type predictability (results from the NHPs, main paper, Fig.4). Please see Methods for MSTD model details. The simulated task was arbitrarily 70±2 time steps long with the reward delivery or withholding of reward at the 55±2 time step. The delivery of reward was simulated with a +1 feedback for 5 time steps to the Reinforcement Learning (RL) agent, whereas the reward withholding was simulated with a feedback of -0.1 for 5 time steps starting at time step 55. The feedback (as mentioned in Methods) was the immediate reward used to calculate the Temporal Difference (TD) error, which was then used to adapt the weights mapping the current state of the RL agent in its environment to an estimated value function. The goal of the RL agent was to learn and predict the value function at any given time point in the task. The reward delivery during rewarding trials and the reward-withholding period during the non-rewarding trials were each considered being individual stimuli for the MSTD model. The number of microstimuli per stimulus was four, with a standard deviation (σ) of 0.1. The decay rate for the memory trace was maintained at 0.985 with the discount factor (*γ*) set at 0.95. The decay rate of the eligibility trace (*λ*) was set at 0.7 with a learning rate (*α*) of 0.7. There were a total of 360 trials in a simulated session.

Two simulations were run, one with chance trial type predictability and another with complete trial type predictability -

Chance trial type predictability: The number of rewarding and non-rewarding trials was equal in the session. The trials were presented randomly. Supp.fig.3 (a) shows that the value function learned under the Microstimulus TD (MSTD) model only differentiates after the reward delivery/withholding period. Such a prediction matches the observed neural response in NHPs S and PG to this task. The value function is incapable of predicting the expected reward given an unpredictable reward landscape in the task until the reward delivery period. Note that in our model the reward itself acts as a stimulus, that is a state, which is represented with its own temporal bases functions and consequently with the corresponding microstimuli (1). The value function is less phasic in its response compared to the TD error that has been shown to predict and model the neural dynamics of deep brain dopaminergic neurons.

Complete trial type predictability: The number of rewarding and non-rewarding trials was equal in a session. The trials were presented in a completely predictable sequence alternating rewarding and non-rewarding trials. The simulation was performed to address the question of how the extended MSTD agent would respond to a completely predictable reward landscape in a trial without a CS. Supp.fig.3 (b) shows that the value function encoded the expected reward in an un-cued task even before the delivery of reward. This prediction of the value function learned under the extended MSTD model was similar to the M1 activity dynamics reported in the un-cued GF task with complete trial predictability in the main text Fig.4.

The occurrence of a stimulus (R, NR, CSR, CSNR) resulted in a trace value of 1 for the corresponding stimulus. The time following the stimulus was coarsely coded as a combination of the decaying trace and the equally spaced temporal bases functions resulting in microstimuli (MS). These MS when multiplied with a weight matrix provided an estimated value of the current state of the trial. We used a slow trace decay rate of 0.985, which allowed the memory traces to linger in time. The weights associated with each state of the environment are updated using the eligibility trace at a learning rate of α.

Therefore, when a NR trial stimulus is activated at the trial’s outcome time, it results in a decaying trace, and by extension decaying NR-microstimuli values. The trace, with a decay rate of 0.985, stretches into the next trial. Therefore, the model continued to remember the occurrence of the NR stimuli and coarsely encodes the time following the NR stimulus in the form of the MS amplitudes. The immediate reward of +1 awarded at the reward time in an R trial is propagated backwards in time, and the weights associated with the “active” MS values are credited, including the NR-MS from the previous trial. Such updates over multiple trials result in an RL agent that can predict the expected value before the delivery of the reward itself in un-cued tasks with complete trial type predictability. This can be thought of as the model being able to remember the time since the NR stimuli and then associate the previous value awarded in the R trial, allowing it to learn, in a predictable situation such as this, that the NR stimuli is followed by a R stimuli.

**Figure 3:**
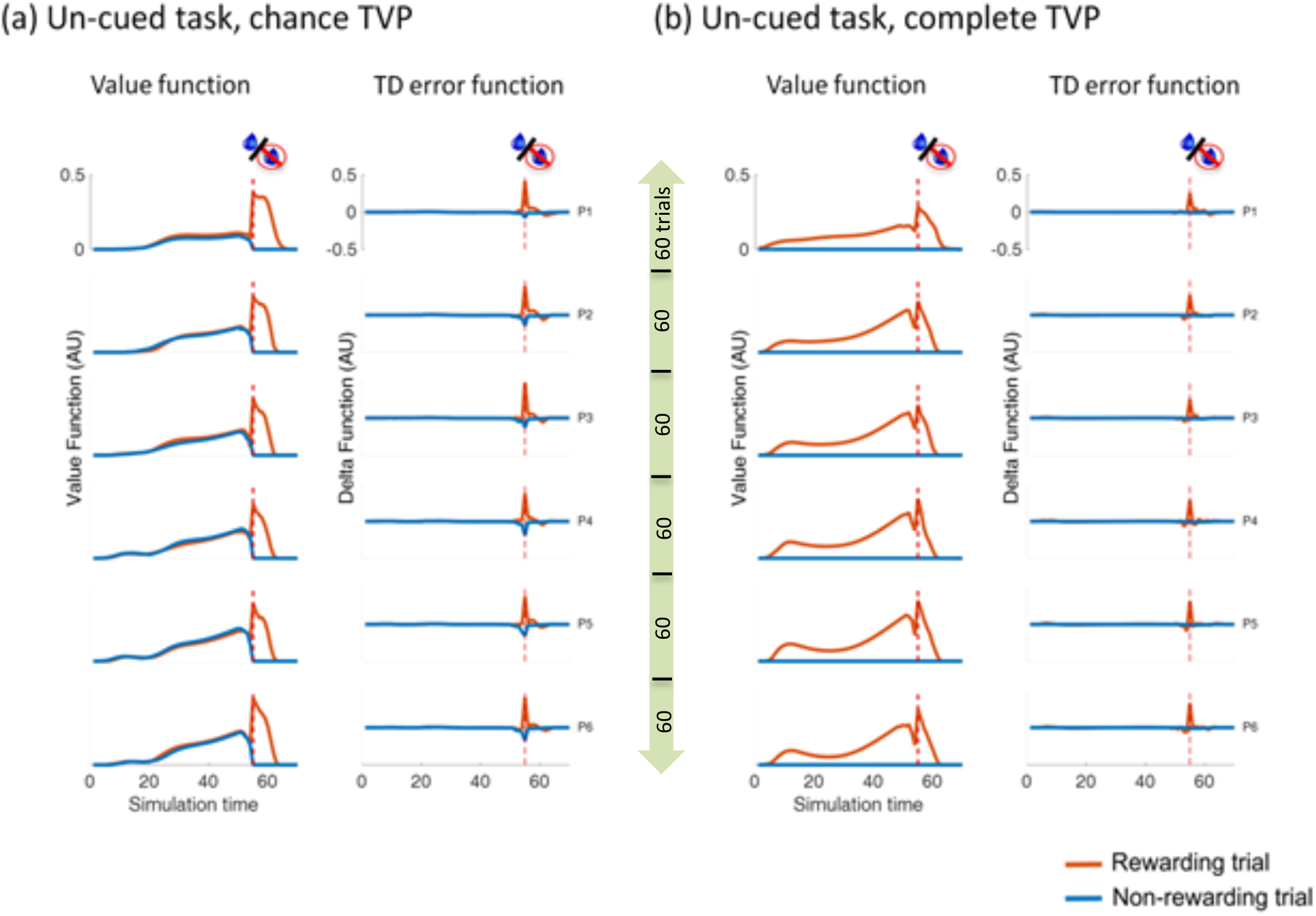
MSTD Value function predicts the expected reward in an un-cued (no CS) task. (a) MSTD simulation of un-cued tasks with chance trial value predictability. The value function and TD error function are presented here for six successive sections (with 60 trials each) that total 360 trials across all sections. The red dotted vertical line represents the reward delivery/withholding period. (b) MSTD simulation of un-cued tasks with complete trial type predictability. The value function and TD error function are presented here for six successive sections of the total number of trials. The red dotted vertical line represents the reward delivery/withholding period.

Cued simulations were also performed with 4 stimuli: the conditioned stimuli CSR and CSNR, as well as the unconditioned stimuli R and NR. Again, the time following a stimulus was coarsely coded with four equally spaced microstimuli. We performed these simulations with chance and complete trial value predictability as with the NHPs. The value functions for R and NR trials differed primarily following the reward delivery in the first few trials of the session. As the session progressed and learning took place, the time steps leading to the reward delivery in R trials were partially credited for the reward, thus leading to an increased expected reward value earlier in the trial itself, regardless of the TVP (Supp.fig.4). Supp.fig.4(b) shows that the value function in sessions with complete trial value predictability captures the expected reward value after conditioning and even before the presentation of the CS, as compared to Supp.fig.4(a). These predictions by the MSTD model are comparable to what were observed empirically in our experiments (Supp.fig.1,2, main paper Fig.3 2).

**Figure 4:**
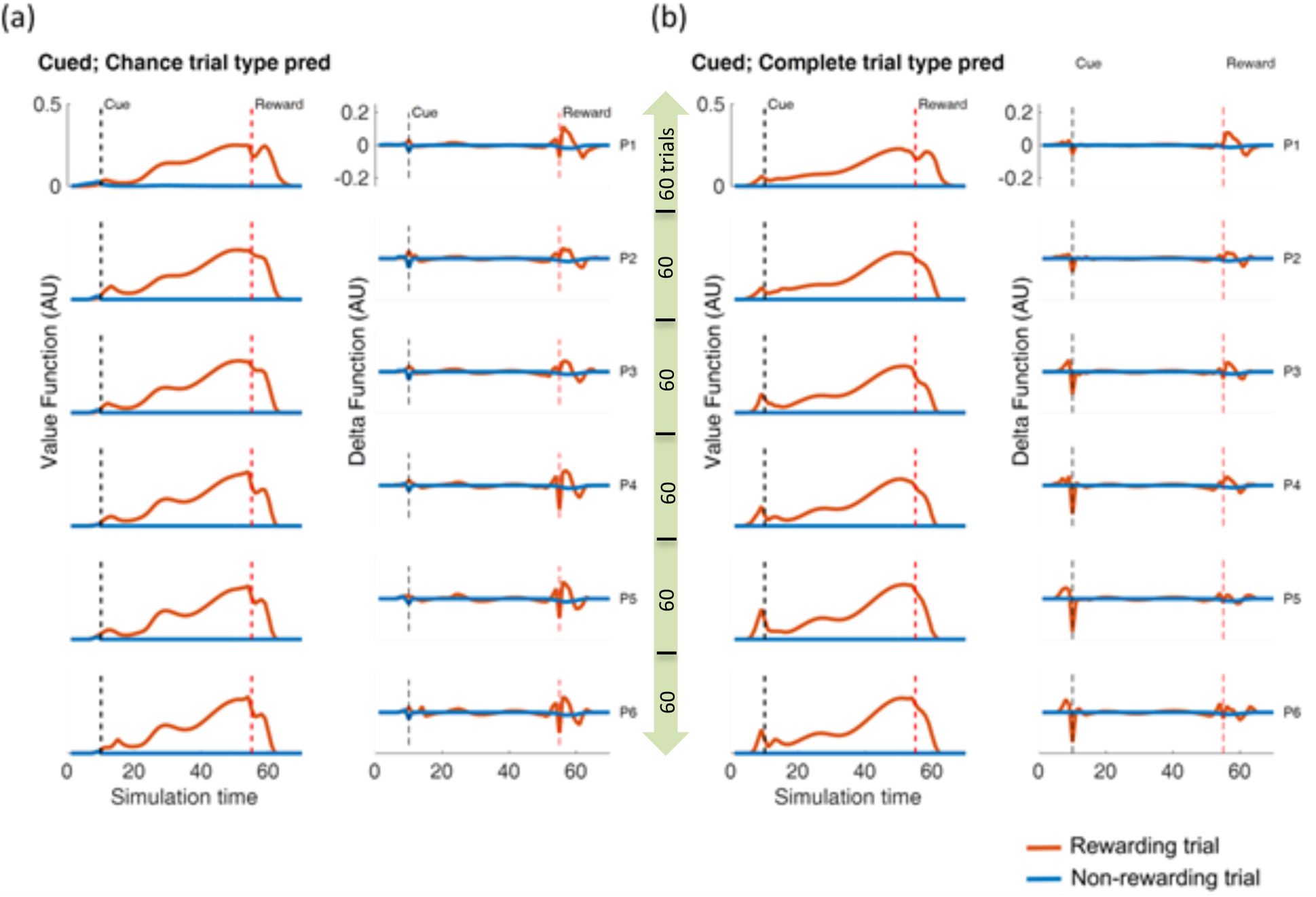
Predictions of the MSTD model in cued trials with varying levels of trial value predictability. (a) Cued trials with chance trial value predictability (b) Cued trials with complete trial value predictability. For both (a) and (b) the first column displays the average value function across R and NR trials in each of the six parts of an evenly divided single session. Each of the six successive sections contains 60 trials that total 360 trials across all sections. The second column captures the temporal difference error (TD error) function across R and NR trials in each of the six parts of a single session. In all subplots the black and red vertical dotted lines indicated the time of cue and the average time of reward, respectively. The time from the cue to reward delivery and the total trial time were centered at 70 and 55, respectively, with a spread ranging from time - 2 to time + 2 time steps.

## References

1. W. Schultz, P. Dayan, R. Montague, A Neural Substrate of Prediction and Reward, Science 275, 1593–1599 (1997).

2. Marsh, V. Tarigoppula, Chen, Francis, Toward an Autonomous Brain Machine Interface: Integrating Sensorimotor Reward Modulation and Reinforcement Learning, Journal of Neuroscience 35, 7374–7387 (2015).

3. Sutton, the ninth annual conference of the … B., A temporal-difference model of classical conditioning, (1987).

4. E. Ludvig, Reinforcement learning in animals, Springer, 2799–2802 (2012).

5. J. O’Doherty, P. Dayan, K. Friston, H. Critchley, R. Dolan, Temporal difference models and reward-related learning in the human brain., Neuron 38, 329–37 (2003).

6. Sutton, Barto, Reinforcement Learning: An Introduction, IEEE Transactions on Neural Networks 9 (1998), doi:10.1109/TNN.1998.712192.

7. Walsh, Anderson, Learning from delayed feedback: neural responses in temporal credit assignment, (2011).

8. J. Hollerman, W. Schultz, Dopamine neurons report an error in the temporal prediction of reward during learning, Nature Neuroscience 1, 304–309 (1998).

9. H. Bayer, P. Glimcher, Midbrain Dopamine Neurons Encode a Quantitative Reward Prediction Error Signal, Neuron 47, 129–141 (2005).

10. G. of the of, Understanding dopamine and reinforcement learning: the dopamine reward prediction error hypothesis, (2011), doi:10.1073/pnas.1014269108.

11. Molina-Luna, Pekanovic, Röhrich, Hertler, Dopamine in motor cortex is necessary for skill learning and synaptic plasticity, (2009).

12. Hosp, Pekanovic, Rioult-Pedotti, Dopaminergic projections from midbrain to primary motor cortex mediate motor skill learning, (2011), doi:10.1523/JNEUROSCI.5411-10.2011.

13. A. Hamid et al., Mesolimbic dopamine signals the value of work, Nature Neuroscience 19, 117–126 (2015).

14. Y. Niv, N. Daw, D. Joel, P. Dayan, Tonic dopamine: opportunity costs and the control of response vigor, Psychopharmacology 191 (2007), doi:10.1007/s00213-006-0502-4.

15. A. Ramakrishnan et al., Cortical neurons multiplex reward-related signals along with sensory and motor information, Proceedings of the National Academy of Sciences 114, E4841–E4850 (2017).

16. Ramkumar, Dekleva, Cooler, Miller, Kording, Premotor and motor cortices encode reward, (2016), doi:10.1371/journal.pone.0160851.

17. McNiel, Choi, and … H., Reward value is encoded in primary somatosensory cortex and can be decoded from neural activity during performance of a psychophysical task, (2016).

18. M. Roesch, C. Olson, Neuronal Activity Related to Anticipated Reward in Frontal Cortex, Annals of the New York Academy of Sciences 1121, 431–446 (2007).

19. H. Cromwell, W. Schultz, Effects of Expectations for Different Reward Magnitudes on Neuronal Activity in Primate Striatum, Journal of Neurophysiology 89, 2823–2838 (2003).

20. E. Ludvig, R. Sutton, J. Kehoe, Stimulus Representation and the Timing of Reward-Prediction Errors in Models of the Dopamine System, Neural Computation 20, 3034–3054 (2008).

21. A. N. McCoy, J. C. Crowley, G. Haghighian, H. L. Dean, M. L. Platt, Saccade reward signals in posterior cingulate cortex, Neuron 40, 1031–40 (2003).

22. D. McNiel, M. Bataineh, J. Choi, J. Hessburg and J. Francis, Classifier Performance in Primary Somatosensory Cortex Towards Implementation of a Reinforcement Learning Based Brain Machine Interface. 32nd Southern Biomedical Engineering Conference (SBEC), pp. 17–18 (2016)

23. Y. Zhao, J. P. Hessburg, J.N.A. Kumar, J.T. Francis, Paradigm Shift in Sensorimotor Control Research and Brain Machine Interface Control: The Influence of Context on Sensorimotor Representations. Front Neurosci. (2018) doi:10.3389/fnins.2018.00579

24. J. An, T. Yadav, J. P. Hessburg, J. T. Francis, Reward Modulates Local Field Potentials, Spiking Activity and Spike-Field Coherence in the Primary Motor Cortex bioRxiv 471151; (2018) doi:https://doi.org/10.1101/471151

25. M. Roesch, C. Olson, Neuronal Activity Related to Reward Value and Motivation in Primate Frontal Cortex, Science (2004)

26. J. H. R. Munsell, Neuronal representations of cognitive state: reward or attention? Trends in Cognitive Sciences, Volume 8, Issue 6, 261 – 265 (2004)

27. Li, Yi et al., Serotonin neurons in the dorsal raphe nucleus encode reward signals, Nature communications vol. 7 10503. 28 Jan. 2016, doi:10.1038/ncomms10503

28. Liu, Zhixiang et al. “Dorsal raphe neurons signal reward through 5-HT and glutamate” Neuron vol. 81, 6 (2014): 1360–1374.

29. Minmin Luo, Yi Li, Weixin Zhong, Do dorsal raphe 5-HT neurons encode “beneficialness”? Neurobiology of Learning and Memory, Volume 135, (2016), Pages 40–49, ISSN 1074-7427

30. Fischer AG, Ullsperger M. An Update on the Role of Serotonin and its Interplay with Dopamine for Reward. Front Hum Neurosci. 2017;11:484. Published 2017 Oct 11. doi:10.3389/fnhum.2017.00484

31. Vertes, R. P. (1991) A PHA-L Analysis of Ascending Projections of the Dorsal Raphe Nucleus in the Rat. The Journal of Comparative Neurology 313, 643–668

32. Pohlmeyer, Mahmoudi, Geng, Brain-machine interface control of a robot arm using actor-critic reinforcement learning. 2012 Annual International Conference of the IEEE Engineering in Medicine and Biology Society, San Diego, CA, 2012, pp. 4108–4111. doi:10.1109/EMBC.2012.6346870

33. Tarigoppula, Rotella, Properties of a temporal difference reinforcement learning brain machine interface driven by a simulated motor cortex. 2012 Annual International Conference of the IEEE Engineering in Medicine and Biology Society, San Diego, CA, 2012, pp. 3284–3287. doi:10.1109/EMBC.2012.6346666

34. Mahmoudi, Pohlmeyer, Prins (2013) Towards autonomous neuroprosthetic control using Hebbian reinforcement learning. J Neural Eng. 2013 Dec;10(6):066005. doi:10.1088/1741-2560/10/6/066005. Epub 2013 Oct 8.

35. Pohlmeyer, Eric A et al. “Using reinforcement learning to provide stable brain-machine interface control despite neural input reorganization” PloS one vol. 9, 1 e87253. 30 Jan. 2014, doi:10.1371/journal.pone.0087253

36. N. Prins, J. C. Snahcez, and A. Prasad, “A Confidence Metric for Using Neurobiological Feedback in Actor-Critic Reinforcement Learning Based Brain-Machine Interfaces,” Frontiers in Neuroscience, 8:111, doi:10.3389/fnins.2014.00111, 2014.

37. Romo, Hernández, Zainos, Brody, Lemus (2000) Sensing without touching: psychophysical performance based on cortical microstimulation.

38. Fitzsimmons, Drake, Hanson (2007) Primate reaching cued by multichannel spatiotemporal cortical microstimulation.

39. Brockmeier, Choi, DiStasio (2011) Optimizing microstimulation using a reinforcement learning framework.

40. Li, Brockmeier, Francis (2011) An adaptive inverse controller for online somatosensory microstimulation optimization.

41. O’Doherty, Lebedev, Ifft, Zhuang (2011) Active tactile exploration enabled by a brain-machine-brain interface.

42. Li (2013) Adaptive inverse control of neural spatiotemporal spike patterns with a reproducing kernel Hilbert space (RKHS) framework.

43. J.S. Choi et. al. Eliciting naturalistic cortical responses with a sensory prosthesis via optimized microstimulation, Journal of Neural Engineering, Volume 13, Number 5 (2016)

44. Chapin, Moxon, Markowitz (1999) Real-time control of a robot arm using simultaneously recorded neurons in the motor cortex.

45. Carmena et al., 2003; Carmena, Lebedev, Crist, O’Doherty (2003) Learning to control a brain–machine interface for reaching and grasping by primates.

46. Hochberg LR, Serruya MD, Friehs GM, Mukand JA (2006) Neuronal ensemble control of prosthetic devices by a human with tetraplegia.

47. Wu W, Gao Y, Bienenstock E, Donoghue J, Black M (2006) Bayesian Population Decoding of Motor Cortical Activity Using a Kalman Filter. Neural Computation 18:80–118.

48. Velliste, Perel, Spalding, Whitford (2008) Cortical control of a prosthetic arm for self-feeding.

49. Gilja V, Nuyujukian P, Chestek CA (2012) A high-performance neural prosthesis enabled by control algorithm design.

50. Chhatbar P, Francis J (2013) Towards a naturalistic brain-machine interface: hybrid torque and position control allows generalization to novel dynamics. PloS one 8:e52286.

51. Chhatbar, von Kraus, Semework, A bio-friendly and economical technique for chronic implantation of multiple microelectrode arrays, (2010).

52. Niv, Daw Dayan, How fast to work: Response vigor, motivation and tonic dopamine, (2005).

53. M. Howe, P. Tierney, S. Sandberg, P. Phillips, A. Graybiel, Prolonged dopamine signaling in striatum signals proximity and value of distant rewards, Nature 500, 575–579 (2013).

54. H. Chase, P. Kumar, S. Eickhoff, A. Dombrovski, Reinforcement learning models and their neural correlates: An activation likelihood estimation meta-analysis, Cognitive, Affective, & Behavioral Neuroscience 15, 435–459 (2015).

55. S. Gershman, Dopamine Ramps Are a Consequence of Reward Prediction Errors, Neural Computation 26, 467–471 (2014).

56. Suri, TD models of reward predictive responses in dopamine neurons, (2002).

57. E. Ludvig, R. Sutton, J. Kehoe, Evaluating the TD model of classical conditioning, Learning & Behavior 40, 305–319 (2012).

## References

1. Ludvig EA, Sutton RS, Kehoe JE (2008) Stimulus Representation and the Timing of Reward-Prediction Errors in Models of the Dopamine System. Neural Computation 20(12):3034–3054.

